# Understanding the molecular mechanism of umami recognition by T1R1-T1R3 using molecular dynamics simulations

**DOI:** 10.1101/597286

**Authors:** Hai Liu, Lin-Tai Da, Yuan Liu

## Abstract

Taste receptor T1R1-T1R3 can be activated by binding to several natural ligands, e.g., L-glutamate and 5’-ribonucleotides etc., thereby stimulating the umami taste. The molecular mechanism of umami recognition at an atomic level, however, remains elusive. Here, using homology modeling, molecular docking and molecular dynamics (MD) simulations, we investigate the effects of five natural umami ligands on the structural dynamics of T1R1-T1R3. Our work identifies the key residues that are directly involved in recognizing the binding ligands. In addition, two adjacent binding sites in T1R1 are determined for substrate binding, and depending on the molecular size and chemical properties of the incoming ligand, one or both these binding sites can be occupied. More interestingly, the binding of varied ligands can lead to either closing or opening of T1R1, based on which, we further classify the five ligands into two groups. This different binding effects are likely associated with the distinct umami signals stimulated by various ligands. This work warrants new experimental assays to further validate the theoretical model and provides guidance to design more effective umami ligands.

**Author summary:** Umami, as the fifth basic taste, is induced by umami substances from the natural food, such as L-glutamate, 5’-ribonucleotides, and peptides etc. These umami substances are widely added to foods as flavor enhancers to promote food quality. However, although extensive experimental and theoretical studies have been devoted to revealing the recognition mechanisms of the taste receptor T1R1-T1R3 to the umami ligands, the detailed molecular mechanism is still unknown, largely due to the lack of the receptor structure. Here, using a new template structure different from the former theoretical studies, we constructed a more accurate homology model of T1R1-T1R3. Based on this receptor model, combined with molecular docking and MD simulations, we investigate how different ligands with varied molecular size and chemical groups might affect the dynamics of T1R1. Our work provides the structural basis for relating the dynamics of umami receptor induced by varied ligands to the resulting umami signal.

## Introduction

The umami, induced by binding of L-glutamic acid salt (MSG) to its taste receptor type 1 (T1R), was discovered as one of the five basic tastes by Ikeda in 1908 [1–3]. T1R is comprised of three members, including T1R1, T1R2, and T1R3, which can further dimerize to form either sweet taste receptor (T1R2-T1R3) or umami taste receptor (T1R1-T1R3) [2, 4]. T1R belongs to the G-protein-coupled receptors (GPCRs) family and is assigned to the same class as the metabotropic glutamate receptor (mGluR) by previous phylogenetic tree analysis [5].

Human T1R1-T1R3 (hT1R1-T1R3) consists of a large extracellular N-Venus Flytrap domain (VFD), and a seven-transmembrane domain (TMD), connected by a small cysteine-rich domain (CRD) (Fig 1) [5]. VFD has been determined to be the ligand binding domain by former site-directed mutagenesis studies using chimeric T1R constructs based on human sweet and umami T1R VFD domains [3, 6], or human and rat T1R VFD domains [7–9]. Notably, apart from VFD, the hT1R1 TMD is also considered as a potential substrate binding site for some umami substances, such as methional [10], and S807 [6]. As a result, the T1R TMD couples with a G protein inside the cell to stimulate the umami signal [11]. More interestingly, the umami can be further enhanced by binding to additional 5’-ribonucleotides, such as disodium 5’-guanylate (GMP) and disodium 5’-inosinate (IMP) [2, 4, 12]. This synergistic enhancement is considered to be the most unique sensory feature of umami compared to other four basic tastes. Notably, in addition to L-Glu, Nelson et al. and Toda et al. found that the hT1R1-T1R3 can also be activated by other standard L-amino acids, whereas hT1R1 are more sensitive to L-Glu compared to other amino acids [2, 8]. Moreover, small peptides, especially those with a molecular weight lower than 3000 Daltons, can also stimulate umami [13–15].

**Fig 1.**
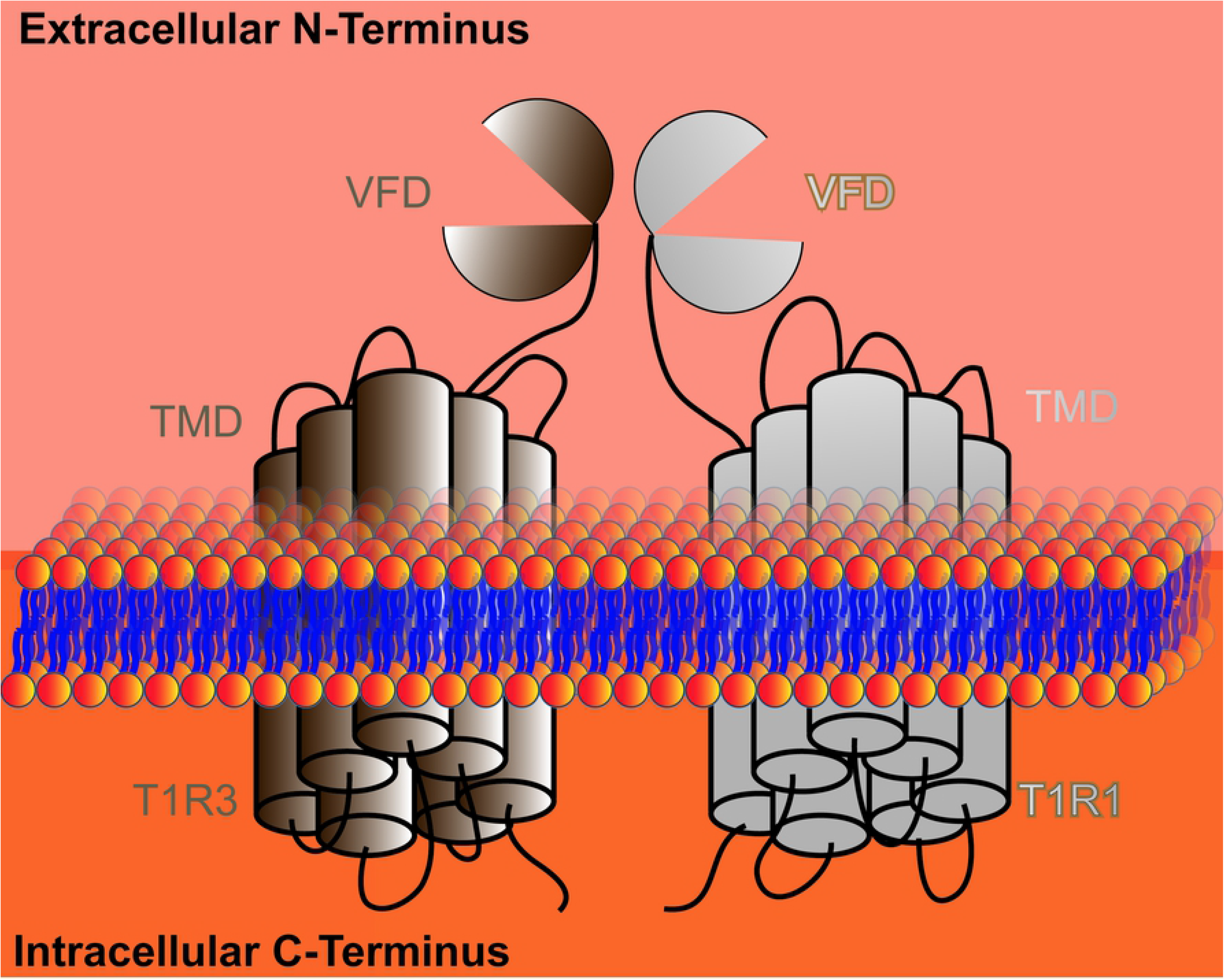
Schematic illustration of T1R1-T1R3 structure, which is a heterodimer with each subunit composed of one TMD and one VFD connected by CRD.

To date, the crystal structure of hT1R1-T1R3 VFD has not been resolved. Computational modeling has been used to construct a homology model of hT1R1-T1R3 and to understand the molecular mechanism of umami recognition. For example, Cascales et al. [16] constructed an hT1R1-T1R3 VFD model using the mGluR1 structure (PDB id: 1EWK) as a template. Their MD simulations reveal that the L-Glu tends to induce the closing of T1R1 but not for T1R3. Likewise, Dang et al. [17] also found that the T1R1 can undergo closing/opening motions in the presence of umami peptides. In addition, Yu et al. [15] constructed a three-dimensional quantitative structure-activity relationship model for five peptides and their umami intensities. Moreover, Dang et al. [18] studied the molecular mechanism of the synergistic effect between L-Glu and umami peptide using molecular docking. They revealed that after T1R1 binds to MSG, T1R3 can adopt an open state that can recruit the peptides, therefore, leading to the synergistic effect on promoting the umami intensity. In sharp contrast, Zhang et al. [7] found that IMP and L-Glu can bind to hT1R1-VFD simultaneously, which provides another explanation for the synergy. In addition, they further identified eight residues that are critical for ligand binding using site-directed mutagenesis. Taken together, the previous theoretical studies have provided some structural basis for the ligand-bound hT1R1-T1R3 complexes. However, the effects of ligand binding on the dynamics of hT1R1-T1R3 are largely unknow, and how structurally varied ligands might affect the dynamics of hT1R1-T1R3 in different manners are also unclear. Moreover, the homology template used in former work is mGluR, which has lower sequence identity (< 30 %) with hT1R1 or hT1R3 compared to the recently obtained crystal structure of fish T1R2a-T1R3 [19].

Here, by employing homology modeling, molecular docking, and MD simulations, we constructed five ligand-bound hT1R1-T1R3 VFD complexes based on the fish T1R2a-T1R3 structure (PDB id: 5X2M), including MSG, WSA (sodium succinate), GMP, IMP, and BMP (beefy meaty peptide). We find all the ligands can reach into a cleft region of hT1R1-VFD, and more interestingly, two binding sites in hT1R1-VFD are identified. Depending on the molecular size and carried charges of the incoming ligands, one or both of the above two sites can be occupied. Moreover, based on the effects of ligand binding on the closing and opening motions of hT1R1-VFD, we further classify the ligands into two groups, which may relate to their different bioactivities as umami substances. Our findings provide an atomistic-level understanding of the structure-function relationship for different substrates, which further guides to design more structurally diversified umami ligands.

## Results

### Structural features of hT1R1-T1R3 VFD in complex with different umami ligands

A homology model of hT1R1-T1R3 VFD was constructed based on the counterpart domain from fish sweet receptor T1R2a-T1R3 (PDB id: 5X2M) complexed with one L-glutamine (L-Gln) ligand. The hT1R1-VFD and hT1R3-VFD subunits share sequence identity of 33.03 % and 33.87 % with the corresponding subunit of fish sweet receptor, respectively, which is higher than mGluR that was used in former studies [3, 8, 17]. Moreover, we constructed the phylogenetic tree for T1R and mGluR based on their VFD, sourced from human, fish, mouse, and rat (S2 Fig). The results show that the hT1R1-T1R3 VFD has higher homology with fish T1R2a-T1R3 compared to mGluR1, indicating that fish T1R2a-T1R3 is a better homology template to construct the hT1R1-T1R3 VFD structure. As shown in Fig 2A, hT1R1-T1R3 VFD is a heterodimer, consisting of two ligand binding (LB) domains that form pseudo-symmetrical lobes, and each LB domain is comprised of two parts, LB1 and LB2, resembling to the structural characteristics of mGluR1 and human GABA(B) receptor [19–21].

**Fig 2.**
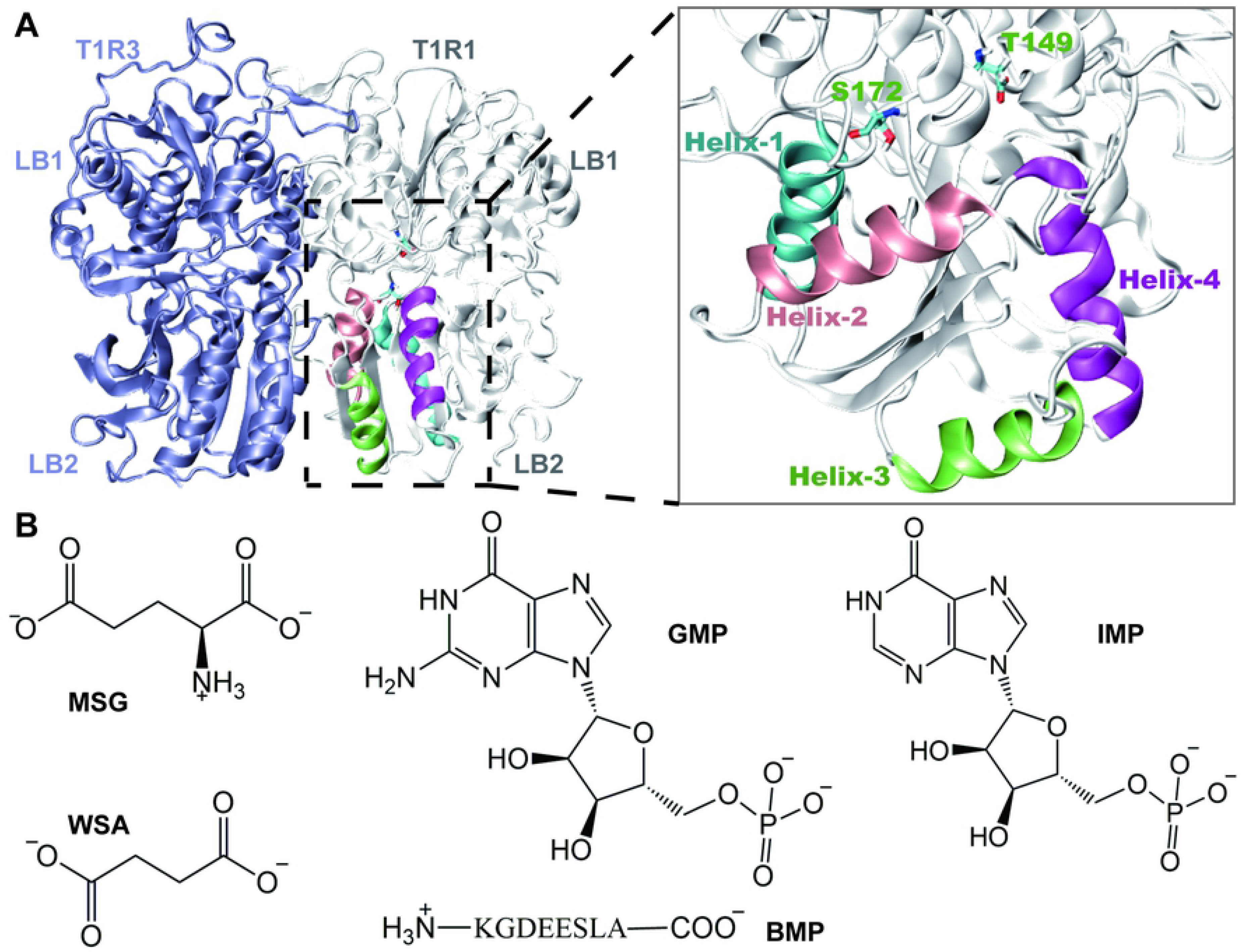
The constructed model of hT1R1-T1R3 VFD and chemical formulas of five umami ligands. **(A)** Homology model of hT1R1-T1R3 VFD, in which hT1R1 and hT1R3 are shown in gray and blue cartoons, respectively. The ligand binding site and key structural motifs of hT1R1 are highlighted in the zoom-in image, in which two active site residues S172 and T149 are shown in cyan sticks, and four LB2 helices, including the residues 192-205 (helix-1), 219-232 (helix-2), 256-266 (helix-3), and 277-290 (helix-4), are highlighted in cyan, pink, lime, and magenta, respectively. **(B)** Chemical structures of five umami ligands studied here, including MSG, WSA, GMP, IMP, and BMP.

Based on the modeled structure of hT1R1-T1R3 VFD, we further constructed ligand-bound hT1R1-T1R3 VFD complexes for five natural umami ligands using molecular docking strategies, including MSG, WSA, GMP, IMP, and BMP (Fig 2B). Only hT1R1 was used as the docking receptor since hT1R3 was found to play an auxiliary role for ligand binding [8, 22]. For each ligand, the structure with the highest binding affinity was used as the final docking complex for the following analysis. In the MSG-bound complex, the ligand can sit in a cleft region formed by LB1 and LB2, and forms direct contacts with the residues S172, T149, A170, D147, A302, R277, and Y200 (Fig 3A). In specific, MSG is sandwiched between the side-chains of D147 and Y220, and forms hydrogen bonds (HBs) with S172 and T149 through its C_α_-COO^−^ group (Figs 2A and 3A). In addition, the C_α_-NH_3_^+^ terminus of MSG can form HBs with the main chain of A170, D147, and A302. The side-chain carboxylate group of MSG can establish one HB with the R277 amide.

**Fig 3.**
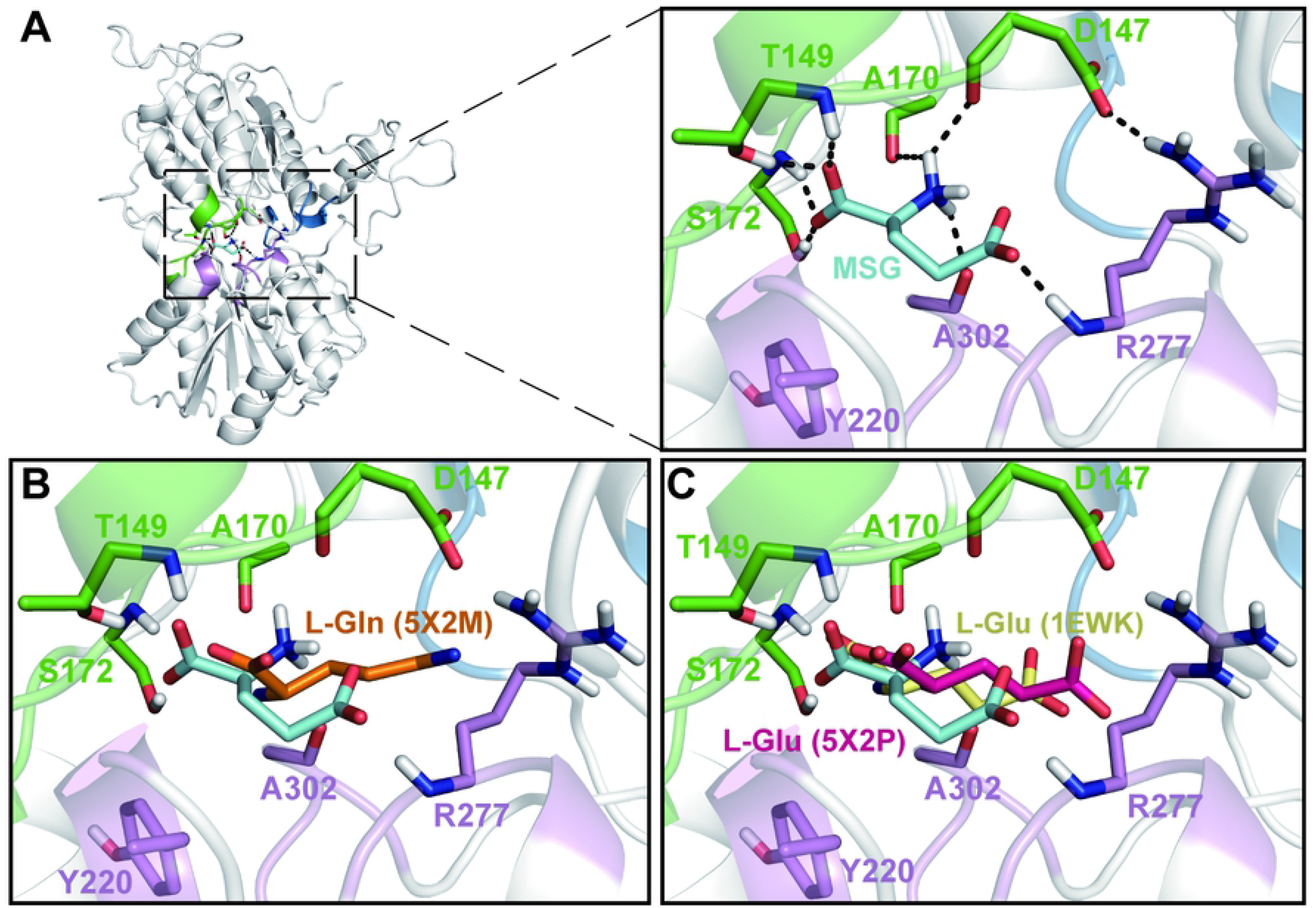
**(A)** Docking complex of MSG-bound hT1R1-VFD. The detailed interactions between MSG and nearby residues are highlighted in the zoom-in image, MSG is shown in cyan sticks and the hT1R1-VFD residues are shown in green/blue/violet sticks. Several key HBs are highlighted with black dashed lines. **(B)** Structural overlay of the MSG-bound hT1R1-VFD to fish T1R2a-T1R3 (PDB id: 5X2M, ligand: L-Gln), and the ligand L-Gln, S150, and T173 in T1R2a-T1R3 are shown in orange sticks. **(C)** Structural overlay of the MSG-bound hT1R1-VFD complex to fish T1R2a-T1R3 (PDB id: 5X2P, ligand: L-Glu) and mGluR1 (PDB id: 1EWK, ligand: L-Glu). L-Glu, S150, and T173 in T1R2a-T1R3 are shown in hot-pink sticks; L-Glu, S165, and T188 in mGluR1 are shown in pale-yellow sticks.

To evaluate the docking result, we compared the MSG-bound hT1R1-T1R3 VFD complex with the crystal structure of fish T1R2a-T1R3 (PDB id: 5X2M, 5X2P) and mGluR1 (PDB id: 1EWK) where the binding ligand is either L-Gln (from 5X2M) or L-Glu (from 5X2P, 1EWK) [19, 20], so that we can compare the ligand binding modes for different complexes. We firstly superimposed our modeled hT1R1-T1R3 VFD to each of the above three crystal structures by their C_α_ atoms. Notably, the C_α_-COO^−^ group of ligands from the four complexes all reside in a same pocket and lie in a similar orientation (Fig 3B and 3C). However, it is worth to note that since the active site residues for different systems are not fully conserved among different systems, the detailed interaction networks between the ligand and nearby residues are also different (S1 Fig). For example, the residue Y220 is completely conserved in the four complexes, while S172 and T149 in hT1R1-T1R3 are replaced by threonine and serine in both T1R2a-T1R3 and mGluR, respectively (Fig 3B and 3C). Moreover, the residue D147 in hT1R1-T1R3 is replaced by glutamine and glycine in fish T1R2a-T1R3 and mGluR, respectively (S1 Fig) [23], which may lead to the fact that the taste stimuli are different for varied species. Taken together, our MSG-bound hT1R1-T1R3 VFD complex provides a fairy good starting structure for the following MD simulations.

We then docked the other four ligands (WSA, GMP, IMP, and BMP) into the same binding pocket as that for MSG. In the WSA binding complex, one of its COO^−^ group sits in the same site as the C_α_-COO^−^ group of MSG, while the other COO^−^ group can form HBs with the residues A302 and S300 (Fig 4A). In addition, WSA can form hydrophobic contacts with Y220 (Fig 4A). For the GMP and IMP binding complexes, the two ligands bind to hT1R1-T1R3 VFD in a very similar mode (Fig 4B and 4C). The nucleobase of these ligands can form hydrophobic contacts with Y220 and several HBs with S172 and T149, while the phosphate group can reach into another pocket and form HBs with S276, S48, N69, and R277. Finally, in the BMP-bound complex, the peptide BMP binds in the same pocket as that for the phosphate group of GMP and IMP (Fig 4B, 4C, and 4D). In specific, the BMP can form HBs with the side-chain of R151 through its C-terminal A8. Moreover, the negatively charged residues D3, E4, E5 of BMP can also form HBs with the residues A302, Y220, S276, R277, S385, and N69 (Fig 4D).

**Fig 4.**
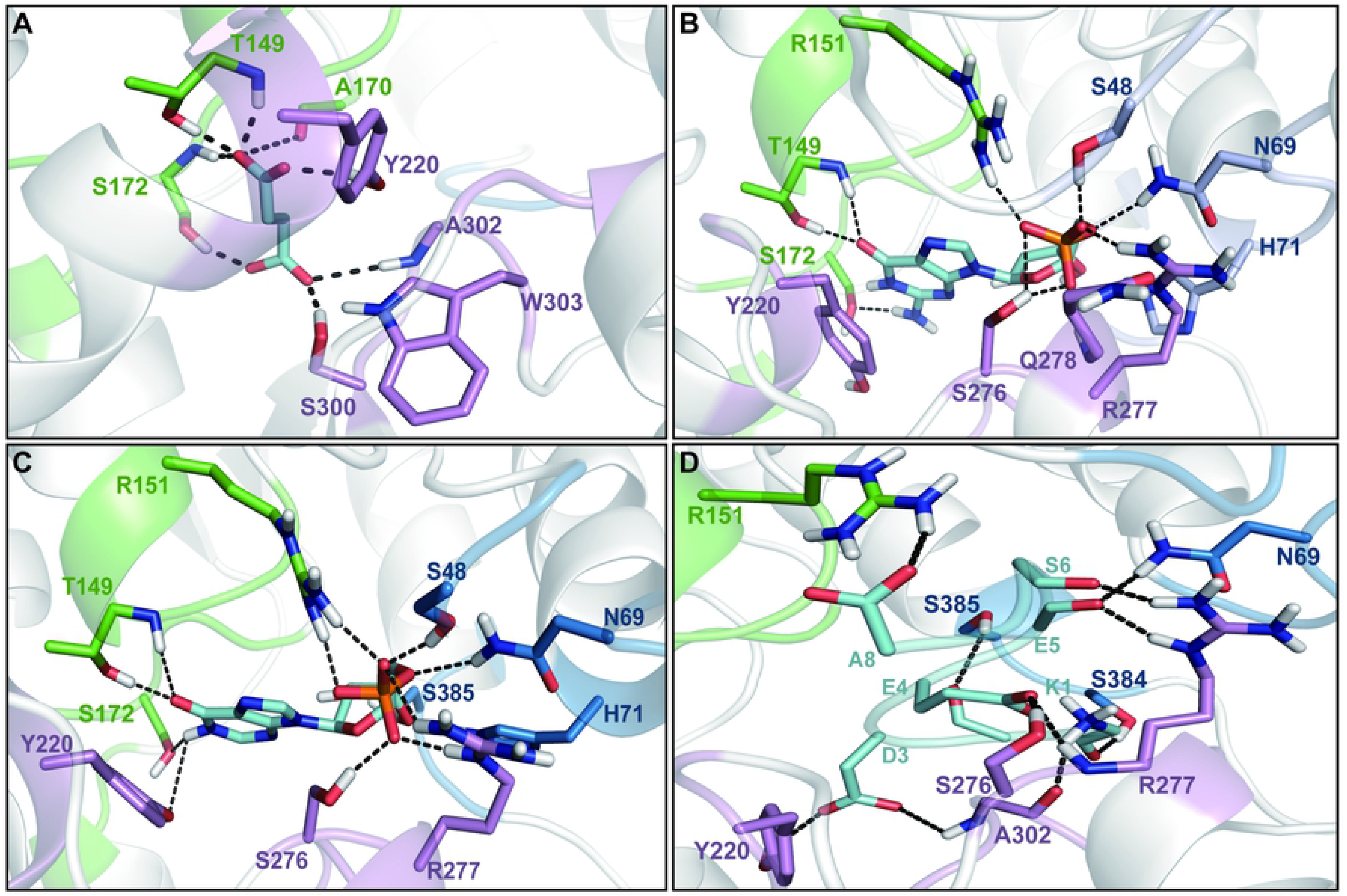
Interaction networks between umami ligands with hT1R1-VFD in each docking complex, including WSA **(A)**, GMP **(B)**, IMP **(C)**, and BMP **(D)**. In each complex, the ligand is shown in cyan sticks, and the key residues that form direct contacts with the ligand are shown in green/blue/violet sticks. Several key HBs are highlighted with black dashed lines.

Taken the above results together, two major binding sites exist in the ligand-binding pocket of hT1R1-VFD, one is surrounded by the LB1 residues D147, T149, R151, A170, and S172 (Figs 3A and 4, shown in green sticks), termed as site A; another consists of the LB1 residues S48, N69, H71, S384, and S385 (Fig 4, shown in blue sticks), termed as site B. In particular, the LB2 residues Y220, S276, R277, Q278, S300, A302, and W303 (Figs 3A and 4, shown in violet sticks) are shared by these two binding sites.

### Effects of ligand binding on structural dynamics of hT1R1-T1R3 VFD

To study the effects of the ligand binding on the structural dynamics of hT1R1-T1R3 VFD, we performed MD simulations starting from the above five ligand-bound hT1R1-T1R3 VFD complexes. For each system, we performed two parallel 100-ns MD simulations initiated from different velocities. As a control, we performed additional 100-ns MD simulations for the apo hT1R1-T1R3 VFD (without ligand). To examine whether the system reaches equilibration within 100-ns MD simulation, we firstly calculated the root-mean-square deviations (RMSD) along the simulation time for each system using the minimized apo hT1R1-T1R3 VFD structure as the reference structure. The results show that the RMSD curves tend to level off after ~80-ns for each system (S3 Fig). We thus only kept the last 20-ns simulation data of each system for further analysis.

For each system, we further calculated the RMSD values for the C_α_ atoms of three different parts of hT1R1-VFD, namely LB1+LB2, LB1, and LB2, by firstly fitting to the corresponding regions to the apo hT1R1-VFD structure. The results show that comparing to LB1, the LB2 and LB1+LB2 domains demonstrates relatively higher structural fluctuations, reflected from their larger RMSD values, which suggests that LB1 keeps relatively rigid compared to LB2 during the simulations (Fig 5A, 5B, and 5C). Therefore, to better characterize the recognition mechanism of these ligands, the LB1 domain of the apo hT1R1-VFD is used as a reference for the further analysis. Moreover, the RMSD calculations indicate that the apo system demonstrates relatively small structural fluctuations compared to the ligand-bound complexes (Fig 5A, 5B, and 5C), suggesting the ligand binding can impose some structural change of hT1R1-VFD. On the other hand, among the ligand-bound systems, the BMP-bound complex exhibits the largest RMSD values for the LB1+LB2 regions, likely due to the relatively bigger molecular size of BMP (Fig 5A). The other ligand-bound systems, however, have similar structural fluctuations for the LB1+LB2 regions (Fig 5A). Notably, the paralleled MD simulations for each system can reproduce the structural features of hT1R1-VFD structure fairly well (Fig 5A, 5B, and 5C), we thus only used one of the two MD simulations for the further analysis.

**Fig 5.**
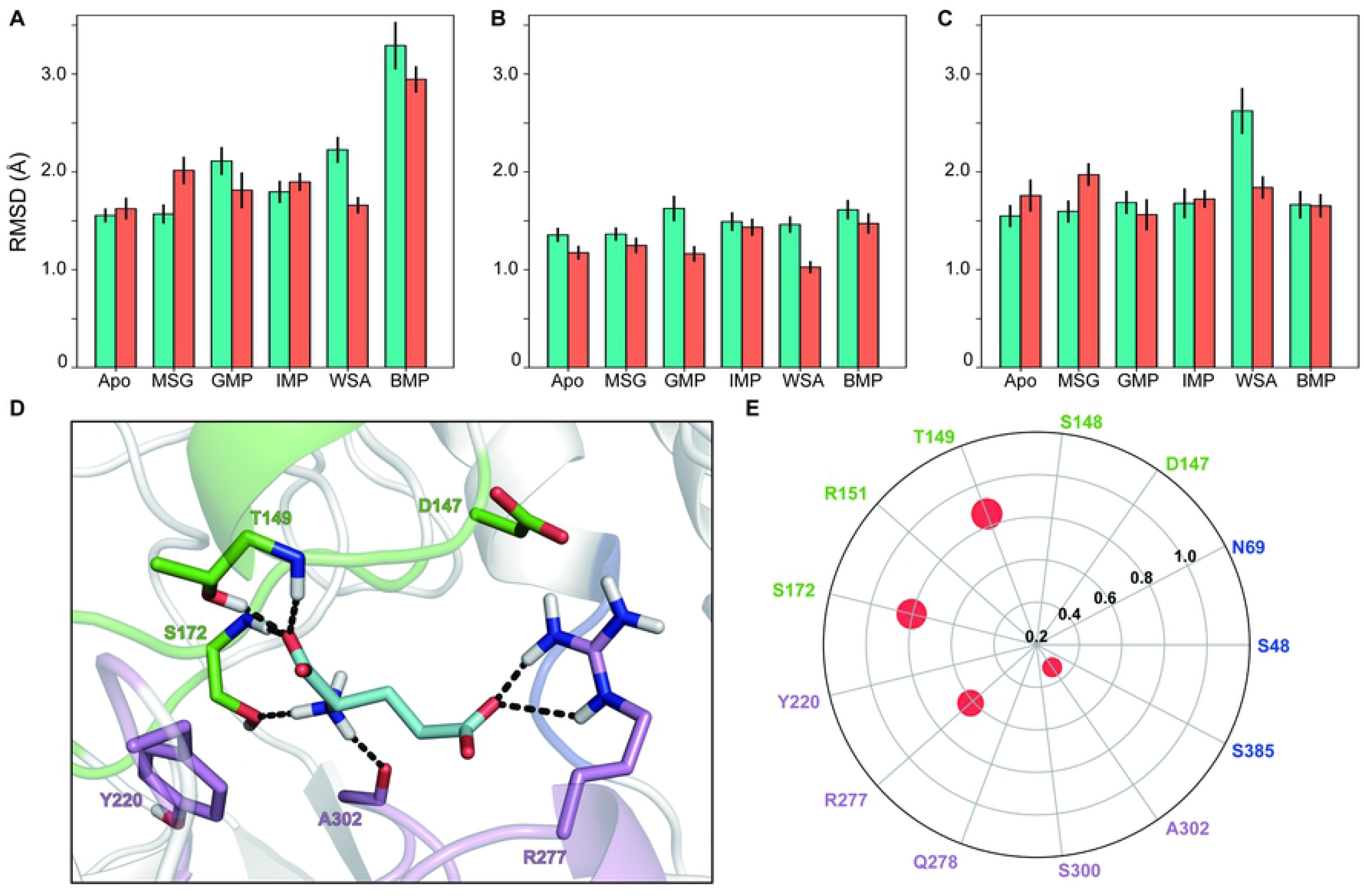
Effects of the ligand bindings on the receptor dynamics and HB network analysis for the MSG-bound hT1R1-VFD complex. RMSD of LB1+LB2 **(A)**, LB1 **(B)**, and LB2 **(C)** regions calculated based on two-parallel 100-ns MD simulations (shown in cyan and coral, respectively). For each system, the mean RMSD and the corresponding standard error were calculated for the last 20-ns simulation conformations. **(D)** The binding mode of MSG (shown in cyan sticks) with hT1R1-VFD, the structure is derived from the last snapshot of the 100-ns MD simulation. The key residues that form direct contacts with MSG are shown in green/blue/violet sticks. Several key HBs are highlighted with black dashed lines. **(E)** The HB network map for the MSG-bound hT1R1-VFD complex. The residues that can form HB with MSG are highlighted with red dot, with the dot size corresponding to the HB occupancy.

We next analyzed the interacting networks between each ligand and hT1R1-VFD by calculating the hydrogen bond occupation (OC_HB_). We only consider the HB that sustains longer than 30 % of the simulation time for further structural analysis. In the MSG system, four residues, namely T149, S172, R277, and A302, can form stable HBs with MSG, among which T149 exhibits the highest OC_HB_ (85.8 %) and A302 exhibits the lowest OC_HB_ (32.9 %) (Fig 5E). In specific, T149 and S172 forms HBs with the C_α_-COO^−^ group of MSG in the binding site A, as observed in the initial docking complex (Fig 3A). Moreover, R277 can form HB with the side-chain COO^−^ group of MSG, and notably, it can also form a salt bridge with D147, which may strengthen the MSG binding (Fig 5D).

Similar HB analyses were performed for other ligand-bound systems. In the WSA-bound complex, four residues, Y220, S300, T149, and S172, can form stable HBs with the COO^−^ groups of WSA, among which Y220 exhibits the highest OCHB (84.5 %) and S172 exhibits the lowest OC_HB_ (38.8 %) (Fig 6A and 6B). In specific, T149 and S172 in the binding site A can form HBs with one of the COO^−^ groups, and the other COO^−^ group can form stable HBs with Y220 and S300 (Fig 6B). It is noteworthy that WSA is unable to bind to the binding site B. In the GMP-bound complex, the ligand can form HBs with T149, A302, and R277, among which T149 exhibits the highest OC_HB_ (99.6 %) and R277 exhibits the lowest OC_HB_ (71.2 %) (Fig 6A and 6C). In the IMP-bound complex, however, six residues, Q278, R151, R277, S172, S48, and D147, form stable HBs with the substrate, and D147 exhibits relatively low OC_HB_ (68.3 %) compared to other five residues (all > 80 %) (Fig 6A and 6D). Notably, GMP and IMP both sit in the same pocket, with their phosphate group forming HB with R277 in the binding site B. Whereas several HBs can be formed between IMP and R151, which is absent in GMP complex (Fig 6C and 6D). Finally, in the BMP-bound complex, BMP largely sits in the binding site B, and forms HBs with S148, S385, R151, and N69, with the OC_HB_ value all higher than 80 % (Fig 6E). The above HBs are formed through the BMP residues E4, E5, and S6. In specific, the side-chain COO^−^ of E4 forms HB with the side-chain of S148, and BMP E5 forms HBs with N69 and S385 (Fig 6E). In addition, BMP S6 can form HB with the side-chain of R151, and an intermolecular HB can also be formed in BMP between K1 and E5.

**Fig 6.**
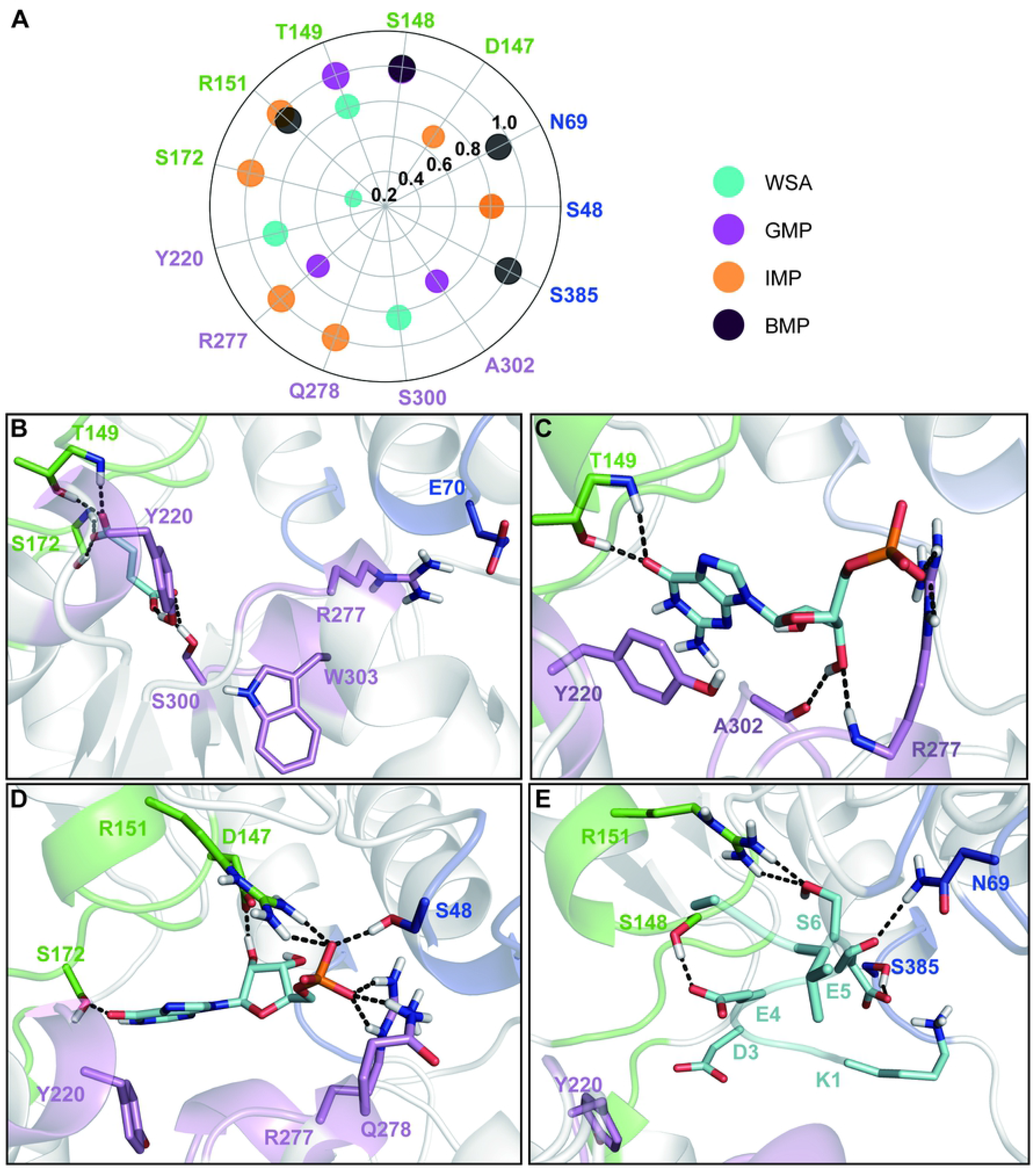
**(A)** The HB network map for four ligand-bound hT1R1-VFD complexes, including WSA, GMP, IMP, and BMP. **(B-E)** The binding mode of each ligand (shown in cyan sticks) with hT1R1-VFD, namely WSA **(B)**, GMP **(C)**, IMP **(D)**, and BMP **(E)**. The structures are derived from the last snapshot of the 100-ns MD simulation for each system. The ligand is shown in cyan sticks, and the residues that form direct contacts with the ligand are shown in green/blue/violet sticks. Several key HBs are highlighted with black dashed lines.

Our HB analysis is consistent with our previous conclusion that, in general, the ligands largely interact with two binding sites in hT1R1-VFD, and in each site, several polar residues within the binding pocket that can potentially form HBs with the incoming ligand, as shown in Figs 5E and 6A. Based on varied molecular size and chemical groups of the incoming ligand, one or both of the above two sites can be occupied, i.e., MSG and WSA largely sit in the site A, while BMP binds to the site B; GMP and IMP, on the other hand, can occupy both binding sites. Therefore, we can expect that varied ligands may impose structural changes of hT1R1-VFD in different manners, which in turn can induce distinct umami intensities.

### Effects of ligand bindings on the opening/closing motions of hT1R1-VFD

Previous studies have suggested that the opening/closing motions of VFD are critical for ligand binding [20]. To measure the effects of the ligands on the opening/closing motions of hT1R1-VFD, we calculated the radius of gyration (Rg) values for the two identified ligand binding sites (A and B) in hT1R1-VFD, which can roughly measure the extents of the hT1R1-VFD opening/closing. Both sites contain the LB2 residues 219-221, 276-278, 300-303, and in site A, we also include the LB1 residues 146-151, 170-173 (shown in green and violet in Fig 7A), and in binding site B, the LB1 residues 47-49, 68-71, 384-386 are included (shown in blue and violet in Fig 7A). For each binding site, we then projected the MD conformations onto two reaction coordinates: Rg and RMSD, calculated for the binding site residues (Fig 7B and 7C).

**Fig 7.**
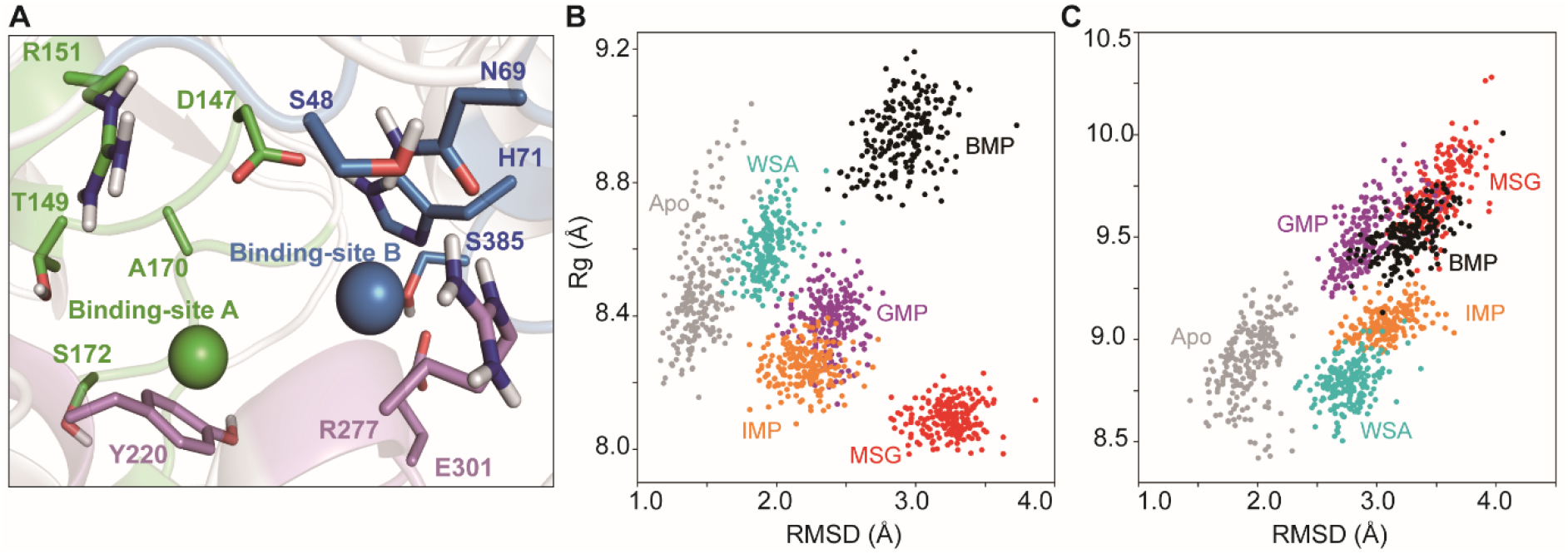
The opening/closing motions of hT1R1-VFD induced by ligand binding. **(A)** To investigate the closing and opening motions of hT1R1-VFD, two binding sites (A and B) are defined and represented with green and blue sphere, respectively. Each sphere represents the centroid of C_α_ atoms of residues within each binding site. In specific, both binding sites include the LB2 residues 219-221, 276-278, 300-303 (shown in violet). In addition, the site A also contains the LB1 residues 146-151, 170-173 (shown in green), and the site B contains the LB1 residues 47-49, 68-71, 384-386 (shown in blue). For the binding site A **(B)** and B **(C)**, we mapped the MD conformations onto two reaction coordinates: one is the Rg of the residues belonging to each site, another one is the corresponding RMSD.

For binding site A, by treating the apo system as a ground state, we can group the five ligands into two classes (Fig 7B). One group includes WSA and BMP that prefer to increase the Rg value of the binding site A, suggesting that they can potentially promote the opening of hT1R1-VFD. In contrast, the other group, consisting of MSG, GMP, and IMP, can tighten the site A, reflected from the reduced Rg values comparing to the apo system. However, it is worth to note that within each group, the ligands also act differently on the protein dynamics, reflected from both the Rg and RMSD calculations. In specific, BMP exerts much higher structural disturbances on hT1R1-VFD than WSA, with an increased Rg and RMSD values, and has stronger effects on promoting the opening of the binding site A compared to WSA. For the second group, MSG tightens the binding site A the most compared to GMP and IMP, reflected from its lowest Rg values (Fig 7B). GMP and IMP, on the other hand, can affect the protein structure in a similar manner.

On the other hand, for binding site B, only WSA can slightly tighten the binding site B, reflected from the reduced Rg value comparing to the apo system (Fig 7C), which is likely due to the fact that WSA is unable to reach the binding site B. The other ligands, however, can all promote the opening of binding site B, reflected from the increased Rg values comparing to the apo system (Fig 7C). In specific, MSG, BMP, and GMP exert relatively larger effects on promoting the hT1R1-VFD opening compared to IMP. It is noteworthy that although MSG largely binds to site A, it tightens binding site A, whereas opens up binding site B (Fig 7B and 7C), we speculate that this is because MSG attracts more interactions from the LB2 residues, e.g., Y220, R277, and A302, which induces the movements of LB2 towards site A.

In conclusion, depending on the various incoming ligands, the hT1R1-VFD can undergo either opening or closing motions. In particular, MSG, GMP, and IMP tend to promote the closing of binding site A, while opening of binding site B. BMP, however, can promote the opening of both two binding sites. Notably, WSA can also promote the opening of binding site A, whereas slightly induce the closing of site B.

### Effects of ligand bindings on hT1R1 LB2 dynamics

The hT1R1-VFD opening/closing induced by ligand binding may result in a structural rearrangement of either LB1 or LB2 domain. Our previous analyses indicate that LB1 keeps relatively rigid compared to LB2, therefore, we further examine the effects of ligand binding on the dynamics of LB2. We thus pinpointed two helices (termed as 1 and 2) near to the binding site A, and two other helices (termed as 3 and 4) near to the binding site B for the following dynamic cross-correlation map (DCCM) analysis (Fig 2A). DCCM describes the correlated motion for a residue pair, with a positive value representing positive correlation (moving towards the same direction), while a negative value indicating the residue pair moving towards the opposite direction (Fig 8).

**Fig 8.**
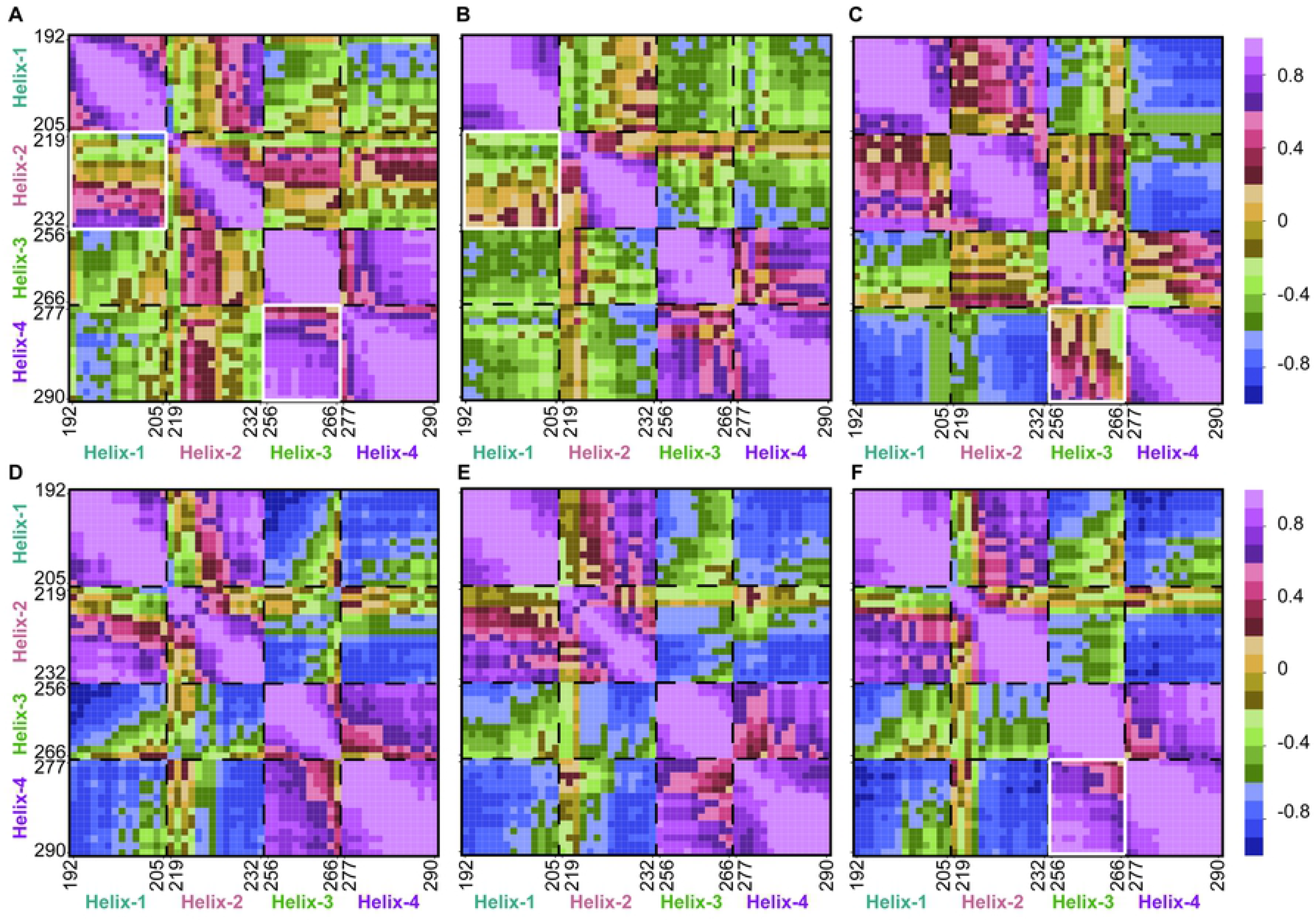
Effects of ligand bindings on hT1R1-VFD dynamics. Cross-correlation matrices of four LB2 helixes for apo hT1R1 **(A)**, and five ligand-bound hT1R1-VFD complexes, namely MSG **(B)**, WSA **(C)**, GMP **(D)**, IMP **(E)**, and BMP **(F)**. The correlation values were calculation based on the last 20-ns MD simulation dataset for each system, with the value equal to 1 representing a positive correlation between two motions, and the value equal to −1 representing a negative correlation between two motions.

The DCCM shows that the five ligands indeed exert different effects on the dynamics of the LB2 domain. For the apo system, helix-1 and 2 have a weak correlation, and helix-3 and 4 have a strong positive correlation (Fig 8A). In comparison, the MSG binding can further weaken the correlation between helix-1 and 2, while has no obvious effect on the helix 3 and 4 (Fig 8B), suggesting that MSG can interfere the interacting networks between helix 1 and 2. In addition, GMP and IMP demonstrate very similar correlation values, suggesting that these two ligands impose similar stresses on the binding sites in hT1R1-VFD, which is consistent with previous results (Fig 8D and 8E). For WSA and BMP, their bindings give rise to a stronger positive correlation between helix 1 and 2 compared to the apo system (Fig 8C and 8F), consistent with the above analysis that they can both open up binding site A (Fig 7B). Moreover, WSA tents to decorrelate helix 3 and 4 compared to all other systems, suggesting its potential role in interrupting the interaction network between helix 3 and 4.

Taken the above results together, the ligand binding within the interface between LB1 and LB2 can indeed affect the structure dynamics of LB2, likely through either tightening or opening up the binding sites. In particular, the closing of binding site tends to weaken the couplings between the adjacent helices, such as the effects of MSG on helix 1 & 2 and WSA on helix 3 & 4.

## Discussion

One of our major findings is to identify the key residues that directly recognize the five umami ligands, which warrants further experimental studies to validate our theoretical models. Former modeling and mutagenesis work has already identified several critical hT1R1-VFD residues for MSG, such as S172, D192, Y220, E301, A170, and A302; moreover, H71, R277, S306, and H308 were identified to be critical for GMP and IMP binding [3, 8]. This result is well consistent with our theoretical models in which S172, A302 can form direct contacts with MSG, and R277 is essential roles in recognizing GMP and IMP. In addition, Y220 can form hydrophobic contacts with all the above three ligands (Figs 5D, 6C, and 6D). Therefore, the constructed model of hT1R1-T1R3 VFD using the fish T1R2a-T1R3 as a template is reliable to derive the binding complexes for umami ligands. Moreover, we can predict more mutational effects on the substrate binding, such as WSA can form important contacts with S172, T149, Y220, and S300 (Fig 6B); the residues T149 and A302 are found to be critical for GMP binding (Fig 6C); five residues R151, S148, N69, Y220, and S385 can directly interact with BMP (Fig 6E). Notably, we can see that the afore-mentioned residues are not highly conserved among different species, such as, S48, N69, D147, R151, A170, Q278, and S385 in hT1R1-VFD are completely non-conservative in fish, rat, and mouse (S1 Fig). Interestingly, former luminescence intensities assay have shown that L-Glu can induce higher umami intensities to hT1R1-T1R3 than mT1R1-T1R3, indicating the sequence differences in the two receptors indeed influences the umami responses [8]. Therefore, our model suggests that these non-conservative residues may contribute to the distinct umami intensities for different species.

More importantly, we find that hT1R1-VFD contains two major binding sites for the umami ligands, and the molecular size and carried charges of the incoming ligand determine its favored binding sites. In specific, MSG and WSA can largely sit in the binding site A, BMP binds to the binding site B, while GMP and IMP can occupy both sites (Figs 5 and 6). From a structural perspective, the site B has a larger binding pocket than site A, which allows the binding of relatively bigger chemical groups (Figs 3A and 7A). On the other hand, to further understand the charge preference for different binding sites, we calculated the electrostatics potentials for hT1R1-VFD using Adaptive Poisson-Boltzmann Solver (APBS). The results indicate that the binding site B is more positively charged compared to binding site A (S4 Fig). In summary, we thus speculate that ligand with large molecular size and negatively charges tends to bind to binding site B, for example, E4 and E5 in BMP, and the phosphate group of GMP and IMP (Fig 6C, 6D, and 6E).

Former experimental studies have shown that the umami threshold value of BMP (1.41 mM) is lower than MSG (1.56 mM), indicating that the taste receptor is more sensitive to BMP than MSG [24]. Moreover, Zhang et al., using the fluorescence imaging plate reader (FLIPR), studied the molecular mechanism of umami synergism for GMP and IMP [3]. They found that both ligands can enhance the umami stimuli of L-Glu. These studies indicate that different umami substances might impose different effects on the protein dynamics. As shown above, different umami ligands can occupy one or both of the two binding sites, leading to distinct closing/opening motions of hT1R1-VFD. For example, BMP promotes the binding site A opening, while MSG tends to close the binding site A. Considering these two ligands have different umami threshold values, we thus speculate the umami properties of the two ligands may relate to their different effects exerted on the protein dynamics. On the other hand, GMP and IMP can impose similar structural influences on the receptor, i.e., they can both induce the closing of binding site A, which lead to a similar dynamic properties of the hT1R1 LB2 region (Fig 8D and 8E). This finding may explain the experimental observation that GMP and IMP have similar umami intensities. Taken together, our work provides a structural basis for relating the effects of ligand binding to the resulting umami signals.

## Conclusions

In this work, by employing homology modeling, molecular docking, and MD simulations, we constructed the ligand-bound hT1R1-T1R3 VFD complexes for five natural umami ligands and revealed how the ligand bindings affect the dynamics of hT1R1-VFD. We identified the key residues that play essential roles in recognizing the ligands, some of the results are supported by former site-directed mutagenesis studies [3, 8]. Further interacting network analysis indicates that two major adjacent binding sites exist in the cleft region of hT1R1-VFD. More interestingly, we find that depending on the molecular size and chemical properties, the incoming ligand can sit in one or both of these two binding sites, which in turn can regulate the closing and opening motions of hT1R1-VFD. In conclusion, our work reveal, at an atomistic-level, the structural basis for the functional roles of different umami ligands in regulating the dynamics of hT1R1-T1R3, and provides further guidance for designing more effective umami ligands.

## Methods

### Constructing the hT1R1-T1R3 VFD structure using homology modeling

By employing the structure of fish taste receptor T1R2a-T1R3 (PDB id: 5X2M) as a homology template, we constructed the model of hT1R1-T1R3 VFD using SWISS-MODEL software [25]. The target amino acid sequences of hT1R1-T1R3 VFD was acquired from the universal protein resource knowledgebase (UniProKB) (HT1R1, Q7RTX1; HT1R3, Q7RTX0). The fish T1R2a-T1R3 was used for the homology modeling because it exhibits the highest sequence identity with hT1R1-T1R3 VFD compared to other homologues (S1 Fig). Moreover, a phylogenetic tree was constructed to further prove that the fish T1R2a-T1R3 is the best template so far for the model construction. In specific, the T1R and mGluR were selected to construct the phylogenetic tree based on their VFD, sourced from human T1R1 (h-T1R1, UniProtKB: Q7RTX1), human T1R2 (h-T1R2, UniProtKB: Q8TE23), human T1R3 (h-T1R3, UniProtKB: Q7RTX0), mouse T1R1 (m-T1R1, UniProtKB: Q99PG6), mouse T1R2 (m-T1R2, UniProtKB: Q925I4), mouse T1R3 (m-T1R3, UniProtKB: Q925D8), rat T1R1 (r-T1R1, UniProtKB: Q9Z0R8), rat T1R2 (r-T1R2, UniProtKB: Q9Z0R7), rat T1R3 (r-T1R3, UniProtKB: Q923K1), fish T1R2a (f-T1R2a, PDBID: 5X2M), fish T1R3 (f-T1R3, PDBID: 5X2M), human mGluR1 (h-mGluR1, UniProtKB: Q13255), human mGluR4 (h-mGluR4, UniProtKB: Q14833), mouse mGluR1 (m-mGluR1, UniProtKB: P97772), mouse mGluR4 (m-GluR4, UniProtKB: Q68EF4), rat mGluR1 (r-mGluR1, UniProtKB: P23385), and rat mGluR4 (r-mGluR4, UniProtKB: P31423).

### Obtaining initial ligand-bound hT1R1-T1R3 VFD complex using molecular docking

By using Gauss View 5 and AMBER 14 software [26, 27], the structure of five umami ligands were constructed, including MSG, WSA, GMP, IMP, and BMP. Then, we performed molecular docking for the above five ligands to hT1R1-VFD using the AutoDock vina-1.1.2 software package with the default scoring function [28]. The docking cube grid box size is set to 30 Å, and the center of grid box is set as the center of mass of L-Gln in the homology template structure. The PDBQT of receptor and ligands was calculated by AutoDock Tools-1.5.6 package [29, 30].

### Setup of MD simulations

Each ligand-bound hT1R1-T1R3 VFD complex was centered in a cubic box with the size of 10 Å, and filled with TIP3P water. A appropriate number of Na+ and Cl^−^ were added in the water box by randomly replacing the solvent waters to neutralize the system and ensure an ionic concentration of 0.15 M. The cutoff distances for Van der Waals and short-range electrostatic interactions were set to 12 Å, and the long-range electrostatic interactions were treated using the Particle-Mesh Ewald (PME) [31]. The SHAKE algorithm was used to constrain all the chemical bonds [32]. The AMBER14SB fore field was employed to describe the hT1R1-T1R3 VFD structure, and the AMBER force fields of ligands were generated using the antechamber module implemented in the Amber-Tools package, and the RESP charges were calculated using the Hartree-Fork methods under the basis set 6-31+g (d) [15, 33, 34].

Energy minimization was firstly performed for each complex by combing the steepest descent and conjugate gradient methods, consisting of the following three steps: (1) with all the heavy atoms of the system constrained with a force constant of 10 kcal/mol/Å^−2^; (2) With all the C_α_ atoms of the system constrained with a force constant of 10 kcal/mol/ Å^−2^; (3) All the atoms are fully relaxed. After minimization, the system temperature was gradually increased from 0 k to 300 k within 100 ps. Then, we performed two 100-ps constrained MD simulations for each complex, with all the heavy atoms constrained with 10 kcal/mol/ Å^−2^ and 5 kcal/mol/ Å^−2^, respectively. Finally, two parallel non-constrained 100-ns NPT MD simulations were conducted under NVT ensemble, each initiated with different velocities. The temperature was kept at 300K using the Langevin Thermostat [35].

### HB and DCCM analysis

One HB was defined as that the distance between that acceptor heavy atom A and a donor heavy atom D is less than 3.5 Å, and the angle formed by atom A, donor H atom and atom D is greater than 135°. The OC_HB_ were determined by the following equation:

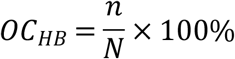

Where n indicates a number of frames that formed HB during MD simulation, N indicates the total number of frames. The PYMOL 2.0 (The PyMOL Molecular Graphics System, Version 2.0 Schrödinger, LLC.) and Visual Molecular Dynamics (VMD 1.9.4a12) [36] were used to analyze and visualize the ligand-bound complex.

DCCM was performed to investigate the correlated motions between four helices from hT1R1 LB2 region. Before constructing the covariance matrix (C_ij_), all the MD configurations were fitted to the reference structure, then C_ij_ is determined by the following equation:

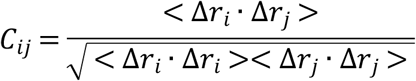

Where the angle bracket represents the average of coordinate over the MD simulation trajectory, and Δr_i_ indicates the deviation of the C_α_ atom of the ith residue from its mean position [37, 38]. The value of C_ij_ fluctuates from −1 to 1. When the value of Cĳ is positive, it represents a positive correlated motion between the ith residue and the jth residue, otherwise, it is negatively correlated motion.

## Supporting information captions

**S1 Fig.**
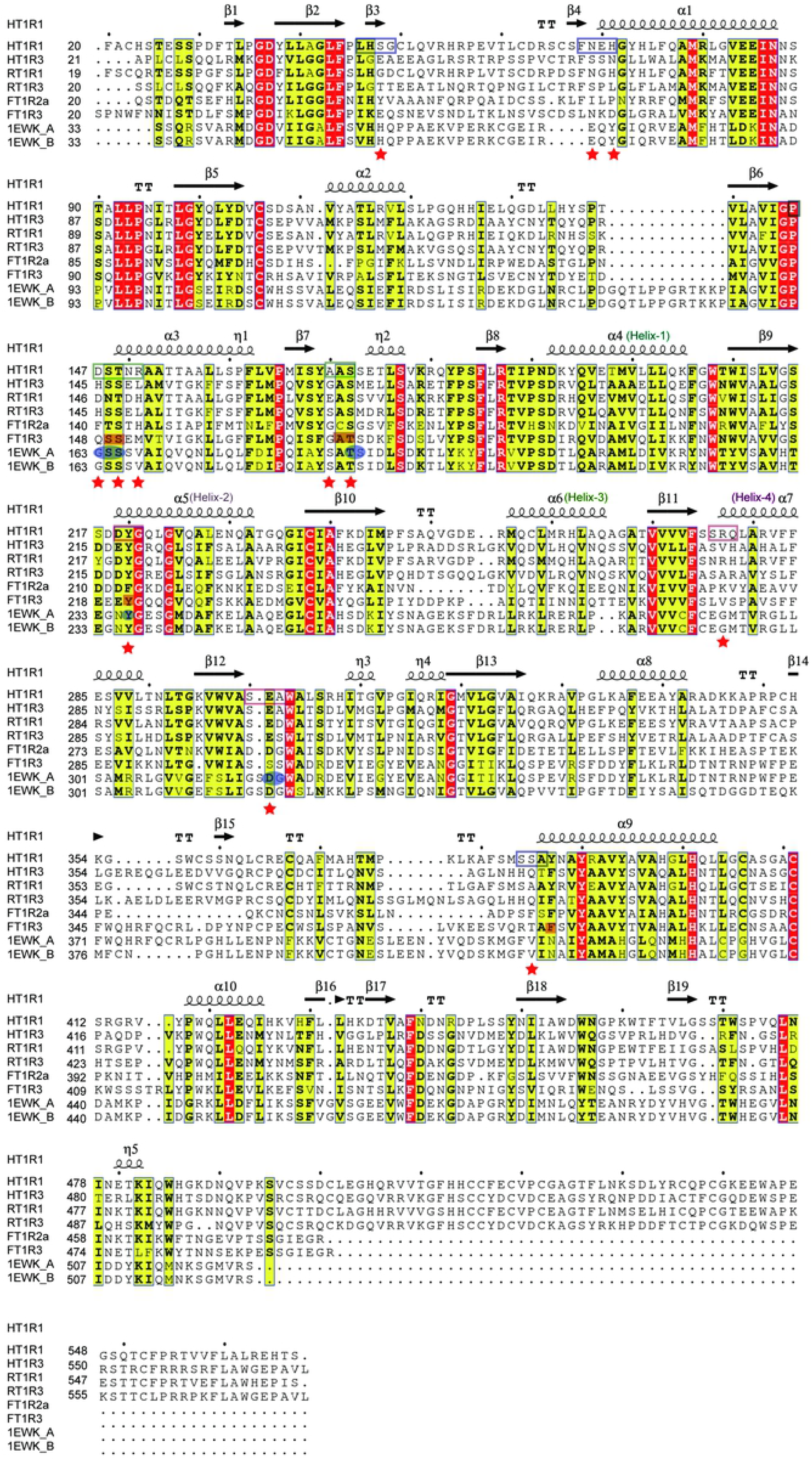
Sequence alignment of the N-Venus Flytrap domain (VFD) in T1R and mGluR1. The amino acid sequences of VFD from human T1R1 (HT1R1, UniProtKB: Q7RTX1), human T1R3 (HT1R3, UniProtKB: Q7RTX0), rat T1R1 (RT1R1, UniProtKB: Q9Z0R8), rat T1R3 (RT1R3, UniProtKB: Q923K1), fish T1R2a-T1R3 (FT1R2a-T1R3, PDBID: 5X2M), and mGluR1 (PDBID: 1EWK) were aligned by blast module of NCBI, and adjusted by ESPript 3.0. The residues 219-221, 276-278, 300-303 are highlighted in violet border; the residues 146-151, 170-173 are highlighted in green border, and the residues 47-49, 68-71, 384-386 are highlighted in blue border. Moreover, the residues that form direct contacts with the five ligand are highlighted with red star. In addition, for FT1R3, the residues that are critical for L-glutamine’s binding are highlighted with coral rectangle. In mGluR1, the residues that play essential roles in recognizing the L-glutamate are highlighted with blue ellipse.

**S2 Fig.**
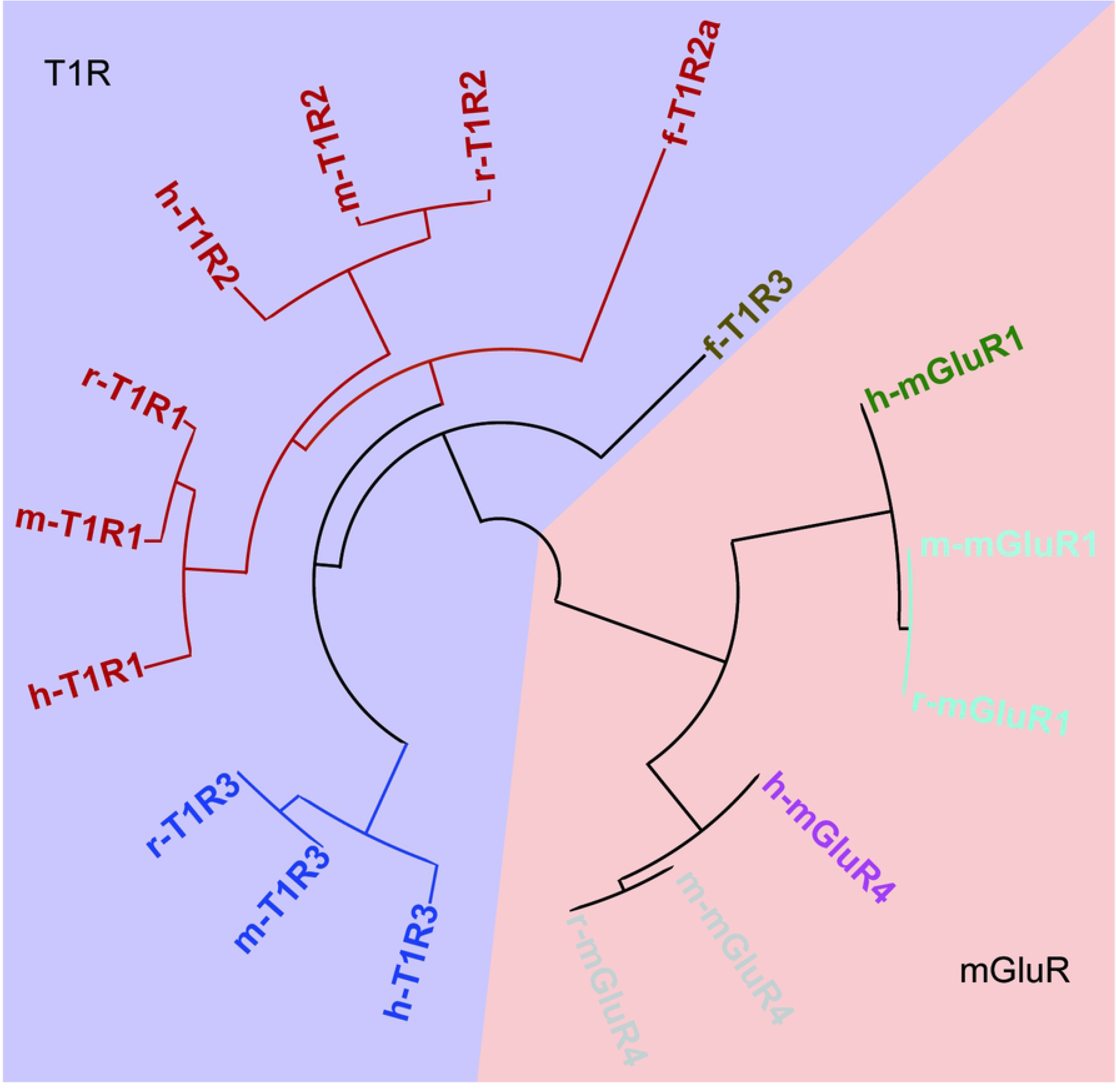
Phylogenetic tree for T1R and mGluR from different species. The phylogenetic tree was constructed using the amino acid sequences of T1R and mGluR VFD, sourced from human T1R1 (h-T1R1, UniProtKB: Q7RTX1), human T1R2 (h-T1R2, UniProtKB: Q8TE23), human T1R3 (h-T1R3, UniProtKB: Q7RTX0), mouse T1R1 (m-T1R1, UniProtKB: Q99PG6), mouse T1R2 (m-T1R2, UniProtKB: Q925I4), mouse T1R3 (m-T1R3, UniProtKB: Q925D8), rat T1R1 (r-T1R1, UniProtKB: Q9Z0R8), rat T1R2 (r-T1R2, UniProtKB: Q9Z0R7), rat T1R3 (r-T1R3, UniProtKB: Q923K1), fish T1R2a (f-T1R2a, PDBID: 5X2M), fish T1R3 (f-T1R3, PDBID: 5X2M), human mGluR1 (h-mGluR1, UniProtKB: Q13255), human mGluR4 (h-mGluR4, UniProtKB: Q14833), mouse mGluR1 (m-mGluR1, UniProtKB: P97772), mouse mGluR4 (m-GluR4, UniProtKB: Q68EF4), rat mGluR1 (r-mGluR1, UniProtKB: P23385), and rat mGluR4 (r-mGluR4, UniProtKB: P31423).

**S3 Fig.**
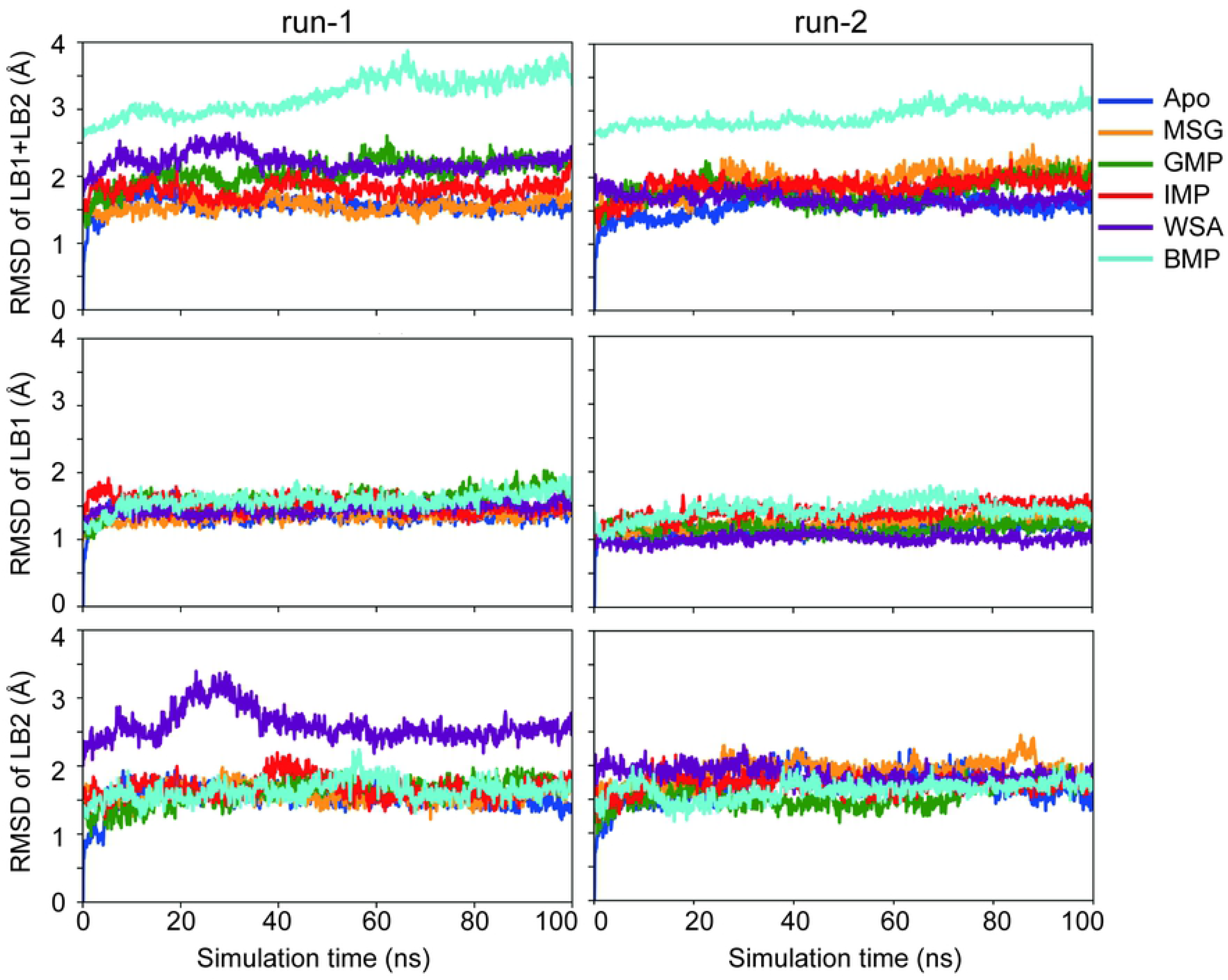
Effects of the ligand bindings on the receptor dynamics. Root mean square deviations (RMSD) of LB1+LB2, LB1, and LB2 regions calculated based on two-parallel 100-ns MD simulations (named run-1 and run-2, respectively). For each system, the RMSD along the simulation time was calculated for the 100-ns simulation dataset.

**S4 Fig.**
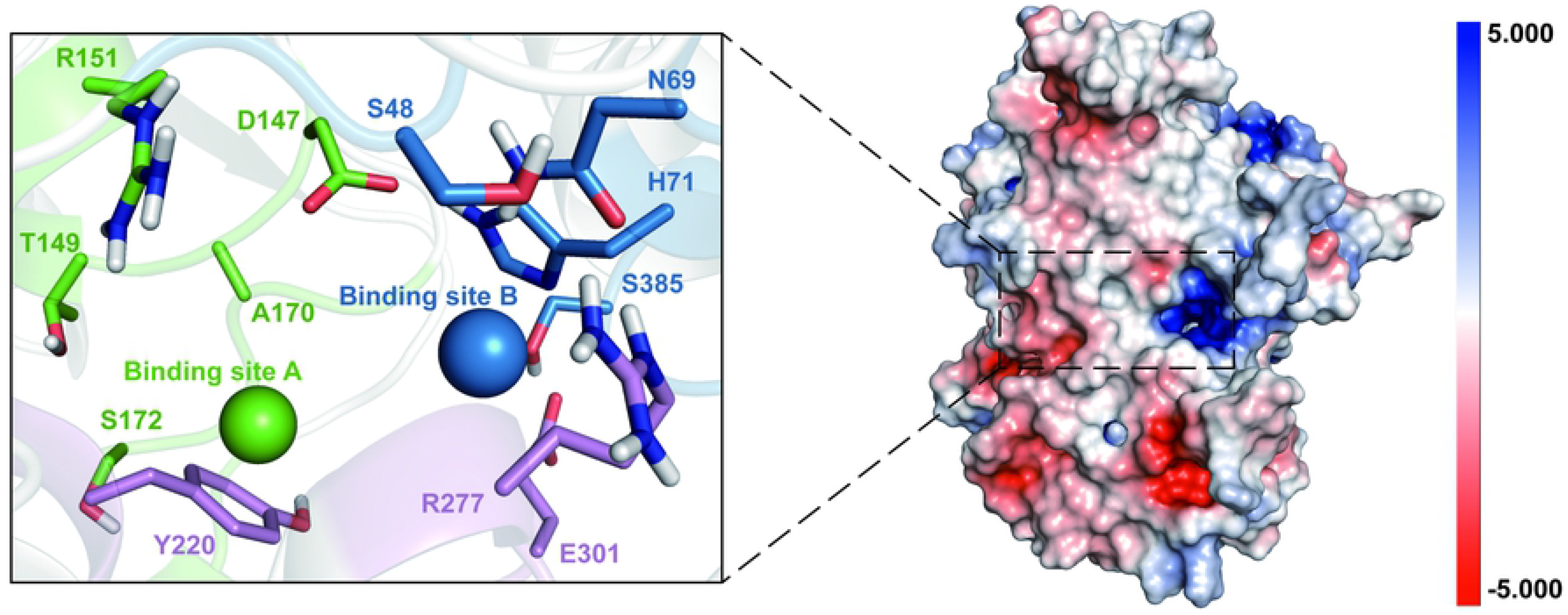
The electrostatics potential of hT1R1-VFD. Positive and negative potential is shown in blue and red, respectively.

## Author Contributions

**Conceptualization:** Hai Liu, Lin-Tai Da, Yuan Liu.

**Data curation:** Hai Liu.

**Formal analysis:** Hai Liu, Lin-Tai Da.

**Funding acquisition:** Yuan Liu.

**Methodology:** Hai Liu, Lin-Tai Da, Yuan Liu.

**Project administration:** Yuan Liu.

**Supervision:** Yuan Liu, Lin-Tai Da.

**Writing - original draft:** Hai Liu.

**Writing – review & editing:** Hai Liu, Lin-Tai Da, Yuan Liu.

